# Flowtigs: safety in flow decompositions for assembly graphs

**DOI:** 10.1101/2023.11.17.567499

**Authors:** Francisco Sena, Eliel Ingervo, Shahbaz Khan, Andrey Prjibelski, Sebastian Schmidt, Alexandru I. Tomescu

## Abstract

A *decomposition* of a network flow is a set of weighted paths whose superposition equals the flow. The problem of characterising and computing safe walks for flow decompositions has so far seen only a partial solution by restricting the flow decomposition to consist of paths, and the graph to be directed and acyclic (*DAG*). However, the problem of decomposing into closed walks in a general graph (allowing cycles) is still open.

In this paper, we give a simple and linear-time-verifiable complete characterisation (*flowtigs*) of walks that are *safe* in such general flow decompositions, i.e. that are subwalks of any possible flow decomposition. Our characterisation generalises over the previous one for DAGs, using a more involved proof of correctness that works around various issues introduced by cycles. We additionally provide an optimal *O*(*mn*)-time algorithm that identifies all maximal flowtigs and represents them inside a compact structure. We also implement this algorithm and show that it is very fast in practice.

On the practical side, we study flowtigs in the use-case of metagenomic assembly. By using the species abundances as flow values of the metagenomic assembly graph, we can model the possible assembly solutions as flow decompositions into weighted closed walks.

Compared to reporting unitigs or maximal safe walks based only on the graph structure (*structural contigs*), reporting flowtigs results in a notably more contiguous assembly. Specifically, on shorter contigs (75-percentile), we get an improvement in assembly contiguity of up to 99% over unitigs, and on the 50-percentile of contiguity we get an improvement of up to 17% over unitigs. These improvements that flowtigs bring over unitigs are 4–14× larger that what structural contigs bring over unitigs.

## 1 Introduction

Network flows are a useful model in assembly problems, since they do not only take into account the graph structure, but also abundance information. In practice, this information is often readily available. For example, in genome or metagenomic assembly, the nodes or the arcs of an assembly graph (e.g. a *de Bruijn graph* [12]) are labelled with the number of times their corresponding string has been observed in the input reads [23, 25, 35, 39]. As another example, in RNA transcript assembly, many tools use *splice graphs* whose nodes (corresponding to exons) and arcs (corresponding to exon junctions) are labelled with their RNA-seq read abundances [11]. Given these abundances, a solution to the assembly problem can be modelled as a *flow decomposition* into weighted paths or walks induced by such abundance values. In the case of perfect data, the superposition of these weighted walks matches the given flow. As a complication, in practice, a flow typically admits a heap of different flow decompositions. This ambiguity is a common issue in the assembly problem (see e.g. [22]), and hence research has focused on reporting only so-called *safe walks* (modelling *contigs* output by modern assemblers), which are partial solutions that are common to all solutions, and hence must also be part of the true DNA or RNA sequence [49].

For splice graphs in RNA transcript assembly (which are in directed acyclic graphs, *DAGs*), Ma et al. [51] gave the first algorithm to decide when a given set of arcs is *safe* for flow decompositions, i.e., when the arcs in the set appear in some, and the same, path of any flow decomposition of the flow in the splice graph. When the arcs in the set form a path, Khan et al. [21] improved the algorithm of Ma et al. [51] from quadratic-time to linear-time, using a simple characterisation of such *safe paths* via a notion of *excess flow* (the flow on the first edge minu the flow leaking from the internal nodes of the path). This also led to an optimal *O*(*mn*)-time algorithm identifying *all* maximal safe paths for flow decompositions in DAGs [21], where *n* and *m* denote the number of nodes and arcs in the graph, respectively. The experimental results on perfect splice graphs from Khan et al. [21] show that safe paths for flow decompositions cover around 18% more of the ground-truth RNA transcripts than paths which are safe based only on the graph structure. Specifically, Khan et al. compare against *extended contigs* [21] (i.e., unitigs extended forwards, as long as nodes have unit out-degree, and backwards, as long as nodes have unit in-degree, also known as *Y-to-V contigs* [18, 22, 30] in the context of genome assembly, where they are close to optimal [49], see Figure 1 (b) for an example). A reason behind the improvement is that the differences between the abundance levels of the different RNA transcripts in a sample can be multiple orders of magnitude [21, 26]. As such, abundant transcripts have a large *excess flow* and can span over branching nodes that would otherwise break a unitig or an extended contig.

**Figure 1:**
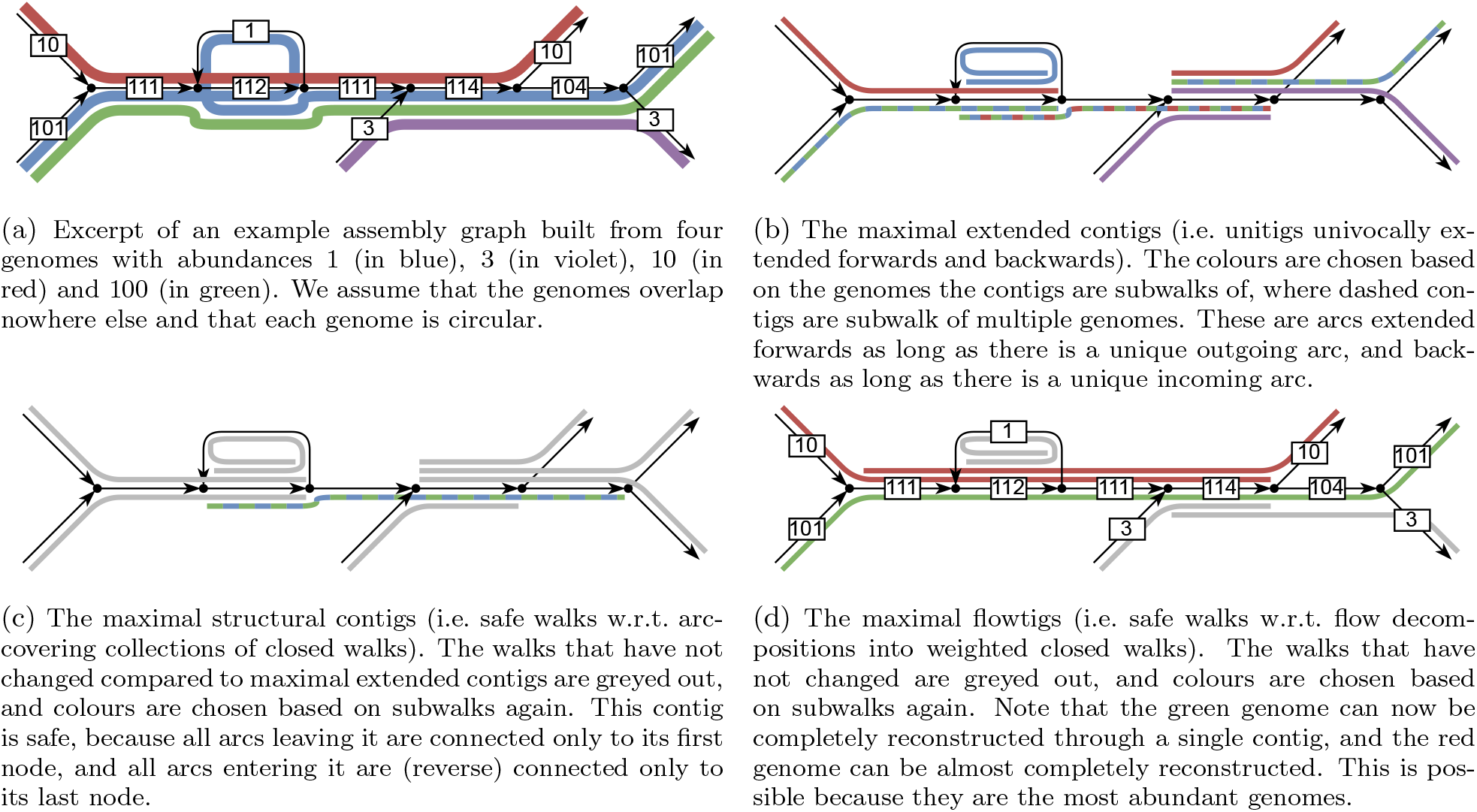
An example assembly graph and various -tigs. All maximal structural contigs are subwalks of flowtigs. Flowtigs produce longer contigs especially for more abundant genomes, as they are not interrupted when meeting a low-abundant genome.

Despite these theoretical and experimental results on safe paths for flow decompositions in DAGs, the analogous theory for non-acyclic assembly graphs is currently lacking. Motivated by the above results for acyclic splice graphs, it is natural to include also abundance information to further restrict the set of assembly solutions in general graphs, to obtain longer safe walks, potentially leading to longer contigs in practice.

### 1.1 Metagenomic assembly and previous safe walks

In this paper we choose metagenomic assembly as the application area to define a concrete notion of genome assembly solution as a flow decomposition, and to evaluate the potential benefits of safe walks for flow decompositions compared to maximal safe walks relative only to the graph structure (*structural contigs*). Metagenomic assembly is crucial for the understanding functionality and composition of various microbiomes containing bacteria, archaea, viruses and single-cell eukaryotes. These microorganism communities are ubiquitous and can be found in multiple ecosystems and multi-cellular organisms, including soil, sea water, the human digestive system, genitals, and others [40]. While there are dedicated metagenomic assemblers for short-read sequencing data (e.g., metaSPAdes [35] and Megahit [25]), there are still distinctive challenges compared to, e.g., single-genome assembly, that are still not completely resolved due to the greatly varying abundances of the species in the sample, and the intra- and inter-species repeats (see e.g. [2]).

With respect to long-read sequencing techniques, the metagenomic assembly problem still raises some challenges, for example when dealing with low abundance levels of high strain variation or bacterial species [50]. Furthermore, the higher cost [8] and larger variance in read length for PacBio HiFi technology [46] as well as the large variance in coverage [23, 46], presents additional issues. While partially alleviated by longer reads, intra-genomic and inter-genomic repeats and inter- and intra-species heterogeneity remain challenging [23,50].

In addition to being an important problem, the metagenomic assembly problem also lends itself nicely to a flow-based theoretical formulation. As in the case of RNA transcripts, the species abundances in a typical metagenomic sample varies by multiple orders of magnitude [27, 42, 47, 48], and this can similarly allow safe walks from more abundant genomes to safely continue over crossings with less abundant genomes. For example, if a branching node has two incoming arcs with abundances 10 and 100, and two outgoing arcs with abundances 90 and 20, then there it is certainly safe to continue from the abundance-100 arc to the abundance-90 arc. See Figure 1 (d) for an example, and compare with Figure 1 (b) and (c) where extended contigs and structural contigs are shorter than flowtigs.

Metagenomic assemblers usually employ overlap graphs [34] or de Bruijn graphs [12], from which they compute unitigs [3, 8, 23, 24, 35] as maximal safe paths and then extend them using possibly unsafe heuristics. Unitigs are non-branching paths in the assembly graph (see Figure 1 (a) for an example). They are *safe*, i.e. they are guaranteed to be subwalks of some genome in the metagenome under the assumption that all genomes are closed walks in the assembly graph such that each arc is covered by some genome, and that the assembly graph is error-free (*arc-centric model*). However, recently, Acosta, Mäkinen and Tomescu [37] have shown that there are longer walks that are safe based on the graph structure, w.r.t. solutions defined as arc-covering collections of closed walks. They have characterised a safe walk *W* by the conjunction of two conditions, and shown that these are both necessary and sufficient. First, there must be no cycle in the graph that contains a subwalk of *W*, but none of its prefixes and suffixes. Such cycles are also called *forbidden paths* [49]. Second, there must be a node *ν* in the graph such that all cycles through *ν* have *W* as subwalk. Acosta, Mäkinen and Tomescu have shown that these maximal structure-based safe walks can be computed in *O*(*m*^2^ + *n*^3^ log *n*) time in node-centric assembly graphs and in *O*(*m*^2^*n* log *n*) time in arc-centric assembly graphs. Later, Cairo et al. [5] have improved this bound for node-centric assembly graphs to *O*(*mn* + *o*), where *o* is the total length of the maximal safe walks.

While these papers close the question of what are the longest walks that can safely be reported from an assembly graph by considering only its structure (assuming perfect coverage), when introducing abundances, the field is much less explored. In the case of a flow decomposition into a single walk where all arcs have abundance 1, i.e. the decomposition into an *Eulerian walk*, the safe walks have been characterised [38]. This model however is not realistic, as the abundances restricted to be only 1 forbid repeats, which are a common structure in genomes. Additionally, a model with restricting the flow decomposition to a single closed walk (but without restricting the abundances to value 1) was considered before [20], but not under the perspective of safety. Such a model however would not be useful for metagenomic assembly, as it assumes just one genome.

In line with previous studies assessing the potential benefits of using longer safe paths or walks [4, 21, 49], in this paper we assume an error-free setting. As such, each circular genome (with its abundance) corresponds to a weighted closed walk in the graph, and thus the superposition of these walks induces a flow where the *flow conservation property* holds at every node, i.e. the sum of incoming flow equals the sum of outgoing flow. Given only the graph and its flow, since we have no further information to decide which flow decomposition is the correct one, a metagenomic assembly is then *any* decomposition of the flow (into weighted closed walks). We study the problem of finding the safe walks in this model, i.e. finding those walks *W* such that for any flow decomposition 𝒟 = {*D*_1_, …, *D*_*k*_} into closed weighted walks, there exists a closed walk *D*_*i*_ such that *W* is a subwalk of *D*_*i*_.

### 1.2 Our contributions

#### Characterisation

On the theoretical side, we provide the first complete characterisation of the safe walks for flow decompositions into weighted closed walks in general graphs via *flowtigs* (walks with positive excess flow). Such complete characterization was formerly only known for flow decompositions into weighted source-to-sink paths in DAGs [21]. Surprisingly, the same characterisation as for DAGs (suitably generalised) still works in general graphs, and it is simpler than the one of structural contigs [5, 37]. Indeed, our characterisation of flowtigs involves only local and simple properties (e.g., a particular set of arcs that interact with the walk in question and their flow values), as opposed to structural contigs, which are characterised by forbidden paths, a more global property related to the existence of a particular type of path between internal nodes of the walk. This makes flowtigs verifiable in linear time in their length and simpler to integrate into real assemblers.

##### Theorem 1 (Safety via flowtigs)

*A walk W is safe for an instance* (*G, f*) *of the flow decomposition problem if and only if it is a flowtig*.

Even though the characterisations are the same, when moving from DAGs to general graphs, the proof of correctness becomes significantly more complicated. Khan et al. [21] prove the unsafety of a non-flowtig *W* by constructing an arbitrary avoiding flow decomposition by taking any leaving arc of *W* . In general graphs, not any leaving arc can be taken, because a wrong decision early on might forces us to traverse *W* in the future, even though it would have been possible to avoid it otherwise. We show how to overcome this by using the leaving arcs of *W* in a particular way, thus allowing us to construct such avoiding decompositions.

#### Enumeration algorithm

Further, we introduce an algorithm that can identify all maximal flowtigs (possibly including duplicates) in *O*(*mn*) time in the worst case, inspired by that of Khan et al. [21]. It first computes a flow decomposition of total size *O*(*mn*) and then identifies the subwalks of the decomposition that are maximal flowtigs. The algorithm then reports the flow decomposition and a list of start and end points of maximal flowtigs within the decomposition. However, the actual time complexity of the algorithm is linear in the size of the flow decomposition, which is often much smaller than quadratic in practice, making it competitive (in terms of runtime) against e.g. computing unitigs or extended contigs. We also give a family of graphs which contain Θ(*mn*) distinct maximal flowtigs, proving that our algorithm is optimal for worst case instances.

##### Theorem 2 (Optimal enumeration of flowtigs)

*Given a flow graph* (*G, f*) *having n vertices and m arcs, all its maximal flowtigs can be identified in O*(||*𝒟*||) ⊆*O*(*mn*) *time and space, where 𝒟 is some flow decomposition of total length at most O*(*mn*). *The time and space bounds are optimal*.

#### Application to metagenomic assembly

On the practical side, we experimentally compare flowtigs against unitigs, extended contigs and structural contigs, on various metagenomic datasets. To focus on the effects introduced by using flowtigs, and in line with previous studies for the DAG case [21, 51], we run our experiments on error-free data. We show that flowtigs provide consistently better assembly contiguity than unitigs on all tested datasets. Moreover, the contiguity improvement that flowtigs bring over unitigs is 4–14 × larger than what extended unitigs or structural contigs provide over unitigs. Our algorithm is very fast also in practice, taking only 2 minutes and using less than 4 GiB of memory to execute on the largest (compacted) graph with 459 thousand nodes and 695 thousand arcs.

We thus hope that flowtigs (i.e. paths/walks with positive excess flow) can be used as a theory-rooted heuristic, both as a standalone technique to compute contigs or to complement current path-finding techniques used by state-of-the-art assemblers. While such applications are outside of the scope of this paper, the two main favourable properties are: (i) Flowtigs are defined by a very simple property, which is also *local* to the nodes and arcs of the flowtig (i.e. excess flow). On the other hand, structural contigs rely on global properties of the graph (the absence of forbidden paths), and thus a single false positive or false negative node or arc can have global effects. (ii) Flowtigs can be less sensitive to local errors than unitigs, extended and structural contigs, because they take abundances into account. In this way, arcs with low abundance and hence on the verge of invalid deletion by an error-correction algorithm can be left untouched for flowtigs, and will affect the resulting contigs only lightly because of their small abundance.

## 2 Preliminaries

### Graphs

A *directed* graph *G* is a tuple (*V, E*), where *V* is the set of *nodes* and *E* the set of *arcs*. We allow graphs to have parallel arcs, that is, two vertices may be connected by more than one arc, but we forbid self-loops (since they can be replaced by a path of length two). As such, for an arc *e* ∈ *E* from *u* to *ν*, we define *t*(*e*) := *u* to be its *tail*, and *h*(*e*) := *ν* to be its *head*. The *in-neighbourhood* of a node *ν* ∈ *V* is the set *N* ^*−*^(*ν*) of its incoming arcs, and the *out-neighbourhood* is the set *N* ^+^(*ν*) of its outgoing arcs.

A *walk W* is a sequence of nodes alternated with arcs *W* := (*ν*_1_, *e*_1_, …, *ν*_*ℓ−*1_, *e*_*ℓ−*1_, *ν*_*ℓ*_) such that *t*(*e*_*i*_) = *ν*_*i*_ and *h*(*e*_*i*_) = *ν*_*i*+1_. We denote by #_*W*_ (*e*) the *multiplicity* of *e* in *W* (i.e. the number, possibly zero, of occurrences of *e* in *W*). A *u-ν walk* is a walk such that *t*(*e*_1_) = *u* and *h*(*e*_*ℓ*_) = *ν*. A *closed walk* is a *ν*-*ν* walk for some *ν* ∈ *V* . A *path* is a walk where all *ν*_*i*_ are unique. For paths, *ν*_*ℓ*_ = *ν*_1_ is allowed, in which case it is a *closed path* (we interchangeably use the term *cycle*). In what follows, we will be usually working with walks along cycles, so we define a *recurring cycle* with respect to a cycle *C* to be a walk beginning with *C* followed by a (possibly empty) prefix of any number of concatenations of *C* with itself. For a walk *W*, we denote by |*W* | its number of its arcs. For a set of walks *𝒲* = {*W*_1_, …, *W*_*k*_} we denote its *total size* (number of arcs) by ||*𝒲*|| = |*W*_1_| + … + |*W*_*k*_|.

If *W*_1_ = (*u*_1_, *e*_1_, …, *e*_*ℓ−*1_, *u*_*ℓ*_) and 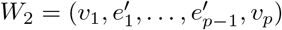are walks such that *u*_*ℓ*_ = *ν*_1_, then 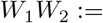 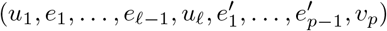 denotes their concatenation. If *e* = (*u*_*ℓ*_, *x*), we analogously define the concatenation of a walk with an arc as *W*_1_*e* := (*u*_1_, *e*_1_, …, *u*_*ℓ−*1_, *e*_*ℓ−*1_, *u*_*ℓ*_, *e, x*). A graph is *strongly connected* if each pair of nodes *u, ν* ∈ *V* it has a *u*-*ν* path.

### Flows

A *flow f* in a graph *G* = (*V, E*) is a function *f* : *E →* ℚ^+^ such that for any node *u* ∈ *V*, its *incoming flow* 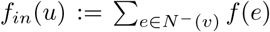 is equal to its *outgoing flow* 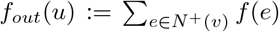 (*flow conservation*). Note that we assume flow conservation *every* node (such flows are also called *circulations*, see e.g., [29, 45]).

A *decomposition 𝒟* of a *flow graph* (*G, f*) is a multiset of *weighted closed walks* (*D*_*i*_, *w*_*i*_), *i* ∈ {1, …, |*𝒟*|} with an associated positive rational weight *w*_*i*_ ∈ ℚ^+^, such that their superposition matches the flow *f*,i.e. 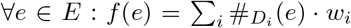 . The *addition f* = *f*_1_ + *f*_2_ of two flows is defined as ⩝ *e* ∈ *E* : *f* (*e*) = *f*_1_(*e*) + *f*_2_(*e*) and the *subtraction f* = *f*_1_ *− f*_2_ is defined as ⩝ *e* ∈ *E* : *f* (*e*) = *f*_1_(*e*) *− f*_2_(*e*). The *multiplication f* = *k* · *f*_1_ of a flow with a scalar *k* ∈ ℚ^+^ is defined as ⩝ *e* ∈ *E* : *f* (*e*) = *k* · *f*_1_(*e*). The *induced flow* of a walk *W* is defined as *f* (*e*) := #_*W*_ (*e*).

The following facts are well known and we refer the reader to e.g. the monographs [1, 45] for further details.

#### Lemma 3.

*If f and f* ^*′*^ *are flows, then f* + *f* ^*′*^ *is a flow and f − f* ^*′*^ *is a flow if it contains only positive νalues. A flow graph* (*G, f*) *has zero or more components all of which are strongly connected*.

### Subwalks and safety

In order to define our safe walks, we first define subwalks of decomposing walks. For a walk *W* that is not closed we define a *subwalk X* to be a walk whose arcs are a substring of the arcs of *W* . For a closed walk *W* we define a *subwalk X* to be a walk whose arcs are a substring of any number of concatenations of *W* with itself. Specifically, *X* may be a substring of only *W*, or *X* starts with a (possibly empty) suffix of *W*, followed by zero or more repetitions of *W*, and ends with a (possibly empty) prefix of *W* . Note that this definition of subwalks of closed walks differs from the usual definition that does not allow *X* to repeat *W* . We use this definition to obtain a more general definition of safety, which results in a more general theoretical result, and in line with previous works such as [37, 49]. We are interested in all the *maximal* safe walks, that is, safe walks such that by extending them with a single arc (in the beginning or the end) renders the walk unsafe.

#### Definition 4 (Safety)

*Let* (*G, f*) *be a flow graph. Then a walk W is* safe *in* (*G, f*), *if for each decomposition 𝒟 into weighted closed walks of* (*G, f*), *it holds that there is some closed walk D* ∈ *𝒟, such that W is a subwalk of D. A* maximal safe walk *is a safe walk that can not be extended without losing the safe property*.

## 3 Characterisation and enumeration of safe walks

We begin by recalling some key notions introduced by Khan et al. [21] for the DAG case. First, we define an arc to be *leaving* from a walk if it is out-going from one of its internal nodes.

### Definition 5 (Leaving arc [21])

*A* leaving arc *of a walk W* = (*ν*_1_, *e*_1_, …, *ν*_*ℓ−*1_, *e*_*ℓ−*1_, *ν*_*ℓ*_) *is an arc e such that ∃i* ∈ {2, …, *ℓ −* 1} : *t*(*e*) = *t*(*e*_*i*_) *and h*(*e*) ≠ *h*(*e*_*i*_).

Next, we recall the notions of *leakage* of a walk (as the total amount of flow that leaves the walk before it ends), and the notion of *excess flow* of a walk (as the flow entering the walk through its first arc, minus its leakage). These concepts will be crucial for showing that a given walk is safe or unsafe. Note that our definition below applies to general graphs, not only to DAGs as in [21]. For example, if there is a leaving arc from a node *ν* for a walk *W*, then its flow value contributes to the leakage of *W* as many times as the number of occurrence of *ν* as internal node of *W* .

### Definition 6 (Leakage and Excess flow [21])

*The* leakage `leakage`(*W*) *and the* excess flow `excess`(*W*) *of a walk W* = (*ν*_1_, *e*_1_, …, *ν*_*ℓ−*1_, *e*_*ℓ−*1_, *ν*_*ℓ*_) *are defined as:*

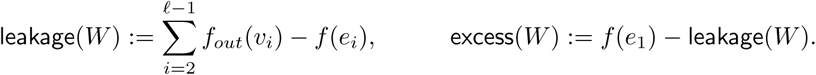

The core idea of the characterisation of safe walks for weighted flow decompositions into source-to-sink paths in DAGs by Khan et al. [21] is to ask where the incoming flow of a given walk can go: certainly, it can flow through the entire walk, but it can also flow through its leaving arcs. The relation between the incoming flow and the leakage is what fully characterises safe walks in DAGs, in fact, safe paths.

For general flow graphs, we define *flowtigs* as those walks with positive excess flow; see Figure 2 for examples.

**Figure 2:**
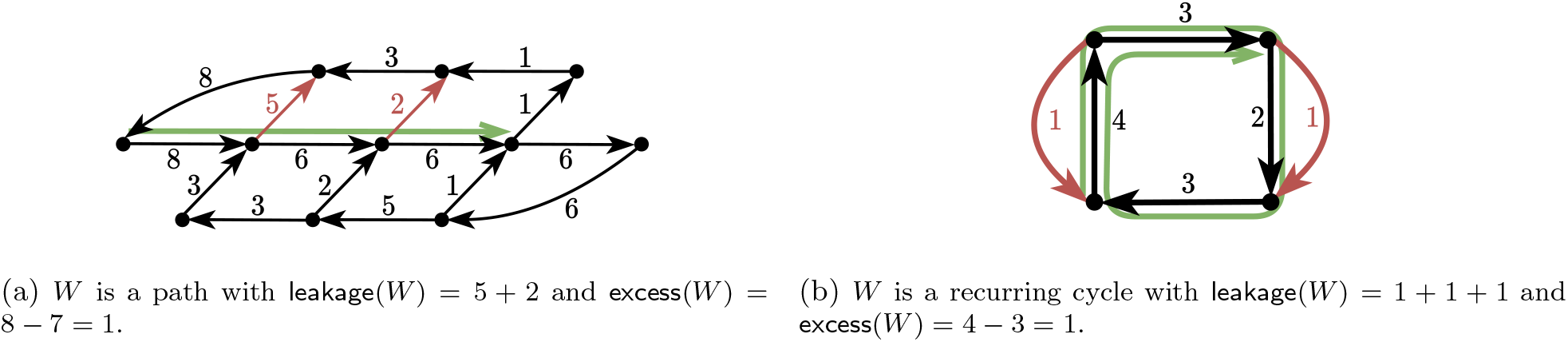
Examples of flowtigs *W* in green with their leaving arcs in red. The numbers denote the flow values of the arcs.

### Definition 7 (Flowtig)

*A walk W is a* flowtig *if* `excess`(*W*) *>* 0.

Our first result below allows us to focus on only two particular types of walks when reasoning about safety, as it imposes restrictions on the shape of any potential safe walk.

### Lemma 8.

*Any cycle followed or preceded by a single arc*

a. *is not safe, and*
b. *its excess flow is non-positive*.

*Proof*. Let *W* = *PC* be a walk consisting of a path made up from a single arc *P* = (*u, e, ν*) followed by a cycle *C* = (*ν, e*_1_, …, *ν*). The case of a cycle followed by an arc is completely symmetric.

For a), let *w* = min_*e∈C*_ *f* (*e*) and let *D* = (*C, w*) be a closed walk (in fact, a cycle) of weight *w*. The walk *W* = *PC* is not a subwalk of any closed walk belonging to a decomposition containing *D*, thus *W* is unsafe.

For b), we first observe that trivially *f* (*e*) *≤ f*_*in*_(*ν*) holds (in general, equality does not hold because other arcs may be entering *ν*). Due to flow conservation, *f*_*in*_(*ν*) units of flow must be carried from *ν* along *C*, and they must eventually exit from *C* (at latest, in *ν*). Note that the nodes of *C* are exactly the internal nodes of the walk *W* . Thus *f*_*in*_(*ν*) *≤* `leakage`(*W*) (again, in general equality does not hold because other arcs may be entering *C*). Thus *f* (*e*) *≤ f*_*in*_(*ν*) *≤* `leakage`(*W*), and thus *W* has non-positive excess flow. □

Consequently, any extension of walks of this form are unsafe, and, therefore, any safe walk is either a recurring cycle or a path (see Figure 2). Importantly, such walks cannot contain leaving arcs of themselves, a fact that we will use in the next results.

### Corollary 9.

*Any safe walk for an instance* (*G, f*) *of the flow decomposition problem into closed walks is either a recurring cycle or a path*.

To prove our characterisation, we will routinely build closed walks that contain a proper prefix of the given walk and then escape via one of its leaving arcs. The following definition addresses this need.

### Definition 10

(Escaping walk). *An* escaping walk *of a walk W* = (*ν*_1_, *e*_1_, …, *ν*_*ℓ−*1_, *e*_*ℓ−*1_, *ν*_*ℓ*_) *is any closed walk containing a non-empty prefix of W, but not containing W as a subwalk*.

We can now show that whenever a walk has positive leakage and does not contain any of its leaving arcs, it admits an escaping walk.

### Lemma 11.

*Let* (*G, f*) *be a flow graph and let W* = (*ν*_1_, *e*_1_, …, *ν*_*ℓ−*1_, *e*_*ℓ−*1_, *ν*_*ℓ*_) *be a walk with* `leakage`(*W*) *>* 0 *such that W does not contain any leaving arc of W itself. Then, for any leaving arc l of W and for any occurrence of t*(*l*) *in W, there is a corresponding escaping walk W*_*l*_ *traversing W until that occurrence of t*(*l*).

*Proof*. Let *l* be any leaving arc of *W* (which must exist since `leakage`(*W*) *>* 0). Let *W*_1_ be the walk along *W* from *ν*_1_ to any occurrence of *t*(*l*) in *W* and let *P*_2_ be a path from *h*(*l*) to *ν*_1_. Such path *P*_2_ always exists by Lemma 3 (note that *t*(*l*) and *ν*_1_ are in the same component). In case *P*_2_ uses any additional leaving arcs of *W*, we let *l* to be the last leaving arc occurring in *P*_2_, and change *P*_2_ to be the path from *h*(*l*) to *ν*_1_. Moreover, for any occurrence of *t*(*l*) in *W*, we can change the prefix *W*_1_ to be until that occurrence of *t*(*l*).

Now, for any *W*_1_ we can define the closed walk *W*_*l*_ := *W*_1_*lP*_2_. Note that *W*_*l*_ does not contain *W* as subwalk because *W*_1_ is a proper prefix of *W* and the only occurrence of *ν*_1_ in *P*_2_ is at the end of *P*_2_, therefore *W*_*l*_ is an escaping walk of *W*. □

We now have everything needed to show our theorem characterising safe walks. To show that a walk *W* with non-positive excess flow is unsafe, we give a constructive argument that builds a flow decomposition where *W* is not a subwalk of any closed walk of the decomposition. We handle separately the two cases from Corollary 9 for a walk to be safe. The idea is to use Lemma 11 which will allow us to escape from *W* in a way that we can control both the decrease in the flow value of the first arc of *W* and in the leakage of *W* .

### Theorem 1 (Safety via flowtigs)

*A walk W is safe for an instance* (*G, f*) *of the flow decomposition problem if and only if it is a flowtig*.

*Proof*. Let *W* := (*ν*_1_, *e*_1_, …, *ν*_*ℓ−*1_, *e*_*ℓ−*1_, *ν*_*ℓ*_). We prove both directions of the statement.

(*⇐*) Suppose *W* is a flowtig, i.e., *f* (*e*_1_) *>* `leakage`(*W*). We prove that *W* is also safe for (*G, f*). Each decomposition of (*G, f*) can carry at most `leakage`(*W*) flow through closed walks that contain *e*_1_ but do not have *W* as subwalk. Therefore, since *f* (*e*_1_) *>* `leakage`(*W*), it follows that each decomposition must contain a closed walk having *W* as a subwalk, so *W* is safe for (*G, f*).

(*⇒*) Suppose *W* is safe, and assume for contradiction that *W* is not a flowtig, i.e. *f* (*e*_1_) *≤* `leakage`(*W*). We argue separately over the two possible cases shown in Corollary 9 for a walk to be safe. In both cases we make use of Lemma 11 to iteratively build a decomposition *𝒟* = {*D*_1_, …, *D*_*k*_} into closed walks where *W* is not a subwalk of any *D*_*i*_.

### Case 1

*W* is a path. As such, it does not contain leaving arcs of itself, and by our assumption, `leakage`(*W*) *>* 0. Thus, Lemma 11 gives an escaping walk *W*_*l*_ (note that the tail of every leaving arc of *W* occurs only once in *W*). 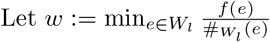 is the largest weight for which the closed walk *W*_l_ with weight *w* “fits” into the flow *f*, and add (*W*_*l*_, *w*) to *𝒟*. In doing so, we update *f* by subtracting the flow induced by *w W*_*l*_ and remove arcs whose flow value becomes zero (so that each *w >* 0), resulting in a new flow by Lemma 3. We repeat this procedure until some arc *e* ∈ *W* disappears from the graph (i.e., its current flow value becomes 0), and then decompose the remaining flow arbitrarily. If this happens, then no walk of the constructed decomposition contains *P* as subwalk, and thus *P* is unsafe for (*G, f*).

To conclude, we argue that if no arc of *W* disappears from the graph before *e*_1_ during our construction, then *e*_1_ will eventually disappear. Since *P* is a path, both *e*_1_ and *t*(*l*) appear only once in *W*_*l*_, and moreover *l* contributes only once in `leakage`(*W*). Then, in each iteration we decrease both *f* (*e*_1_) and `leakage`(*W*) by exactly *w*, and since initially *f* (*e*_1_) *≤* `leakage`(*W*), *f* (*e*_1_) will eventually become zero.

### Case 2

*W* is a recurring cycle. The main difference with respect to the previous case is that a leaving arc *l* of *W* may contribute in `leakage`(*W*) multiple times due to the fact that *W* is a recurring cycle. In fact, if *t*(*l*) occurs *q* times as an internal node of *W*, then *l* contributes to the leakage of *W* exactly *q* times. Then, when subtracting the flow induced by *w* · *W*_*l*_, the leakage of *W* will decrease exactly by *q* · *w* independently of how long the prefix of *W* is in *W*_*l*_ (recall that *W*_*l*_ contains only one leaving arc of *W*), whereas the flow of *e*_1_ will decrease only *p* · *w* times, where *p* can be at most the number of times that *t*(*e*_1_) occurs in *W* . So, even if we build our walk *W*_*l*_ using only one leaving arc of *W*, we may still end up by consuming more from `leakage`(*W*) than from the flow in *e*_1_, possibly making the walk have positive excess flow (see Figure 3c for an example of a decomposition built wrongly in this manner). To overcome this, we will escape *W* via the *last* occurrence of a leaving arc of *W*, which will ensure that *f* (*e*_1_) and `leakage`(*W*) decrease equally.

**Figure 3:**
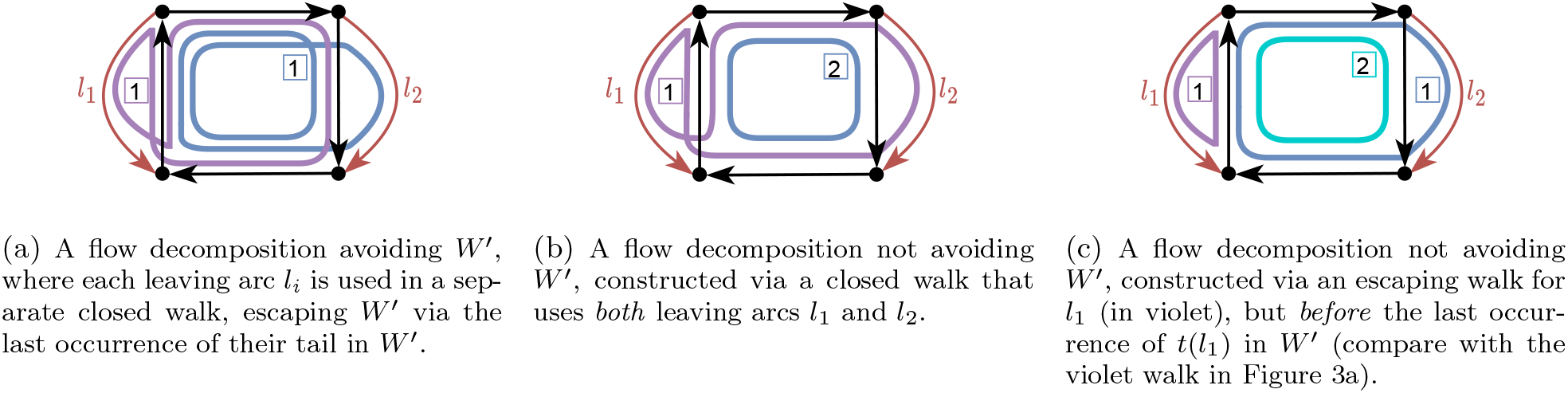
Consider the flow graph from Figure 2b and consider the unsafe recurring cycle *W* ^*’*^ obtained by extending the green (safe) walk *W* forward with the arc of flow value 2. Here we show three flow decompositions of the graph, whose walks are in blue, violet and cyan; their corresponding weights are represented by the boxed numbers.

Since *W* does not contain any of its leaving arcs and `leakage`(*W*) *>* 0, we can apply Lemma 11. As such, we obtain an escaping walk *W*_*l*_, which we require to traverse *W* until the *last* occurrence of *t*(*l*) in *W* . We proceed as in Case 1 and analogously let 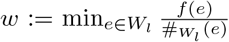, and add (*W_l_ , w*) to 𝒟 until some arc of *W* disappears from the graph (and then decompose the remaining flow arbitrarily).

We again show that during our construction if no arc of *W* disappears from the graph before *e*_1_, then *e*_1_ will eventually disappear. Note that in each iteration we decrease both *f* (*e*_1_) and `leakage`(*W*) exactly by 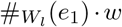, because 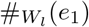equals the number of times *t*(*l*) occurs in *W* (since we took the last occurrence of *t*(*l*) ∈ *W*) which also equals the number of times *l* contributes in `leakage`(*W*), and finally *W*_*l*_ contains no other leaving arc of *W* . Since initially *f* (*e*_1_) *≤* `leakage`(*W*), and we are decreasing some positive value from `leakage`(*W*) at every step (since all *w*’s are positive), *f* (*e*_1_) will eventually become zero. □

We use Theorem 1 to obtain an enumeration algorithm via a standard method, also used by e.g. Khan et al. [21] for the DAG case: first construct an arbitrary flow decomposition into weighted closed walks (which we can do in *O*(*mn*) time and space), and then use a two-pointer approach to check which subwalks of the decomposition walks are maximal flowtigs (since all safe walks are subwalks in any flow decomposition). However, since a flowtig can also loop around a cycle a number of times depending on the flow values (in the worst case), we additionally show that we can avoid this pseudo-polynomial time complexity with an argument related to the ratio between the flow value of the first arc and the leakage of the cycle. As such, we obtain the following theorem:

#### Theorem 2 (Optimal enumeration of flowtigs)

*Given a flow graph* (*G, f*) *having n vertices and m arcs, all its maximal flowtigs can be identified in O*(||𝒟||)⊆ *O*(*mn*) *time and space, where* 𝒟 *is some flow decomposition of total length at most O*(*mn*). *The time and space bounds are optimal*.

See Appendix A for the full proof of this theorem. Therein, we show a family of graphs containing Θ(*mn*) distinct maximal flowtigs, which allows us to conclude that that our algorithm is worst-case optimal. Finally, our algorithm may produce duplicate maximal flowtigs. Therein we also show how to address this using a suffix tree with suffix links, similarly to the approach of Obscura et al. [37]. This increases the runtime of our algorithm to *O*(|𝒟|| log *m*).

## 4 Experiments

We compare flowtigs against unitigs, extended contigs and structural contigs on various datasets. See Appendix A.1 for details about the implementation. As this work primarily focuses on improving assembly contiguity, for storage and deduplication of the assembly sequences we exploited standard Rust libraries, which may not be optimal in terms memory consumption. Although the choice of such string processing methods has no effect on the assembly contiguity, for a more efficient practical implementation one may use more optimised string algorithms and libraries.

Since flowtigs overlap, we cannot use standard metrics for contiguity such as NGA50, because they get artificially inflated through overlaps. Instead, we use the EA50 family of metrics, which is an improvement over NGA50 that is robust against overlapping contigs [44]. The EA50 is computed by aligning the contigs to the reference, and for each reference base identifying the longest contig that aligns to it. These lengths are then sorted and for e.g. EA50, the 50-percentile is reported. EA75 works analogously, by reporting the 75-percentile of largest values. For non-overlapping contigs NGA*x* is equal to EA*x* for all *x*.

### Datasets

See Table 1 for an overview of the metagenomic datasets we use for our experiments. The datasets *simple7, medium20* and *complex32* are from Shafranskaya et al. [47], containing 7, 20 and 32 bacterial genomes, across 6, 19 and 26 genera, respectively. We selected these datasets to be able to see the effects of flowtigs on bacterial communities with various complexities. We refer the reader to [47] for more details on these three datasets. Additionally, we also include the mock community *JGI* [48] which contains 23 bacterial and 3 archaeal strains with finished genomes, which we select as a microbiome that does not only contain bacteria. For more realistic datasets, we select *HMP* [27], which contains 23 high-quality assembled genomes of a real waste water sample. Since the datasets JGI and HMP are missing abundance profiles, we simulate them using the log-normal distribution. This is used by various state-of-the-art metagenomic read simulators [9, 10]. We parameterise the log-normal distribution with a mean of 0 and a standard deviation of 2, which results in a realistic abundance profile for e.g. the human gut [42]. To the best of our knowledge, JGI and HMP are not motivated by specific real metagenomes with available abundance profiles, hence we use this parameterisation to simulate their abundance profiles. To get integer abundances, we round the output of the log-normal distribution up. And for reproducibility, we fix the seed for the random generator. The datasets used for our experiments are available on Zenodo [15].

**Table 1:**
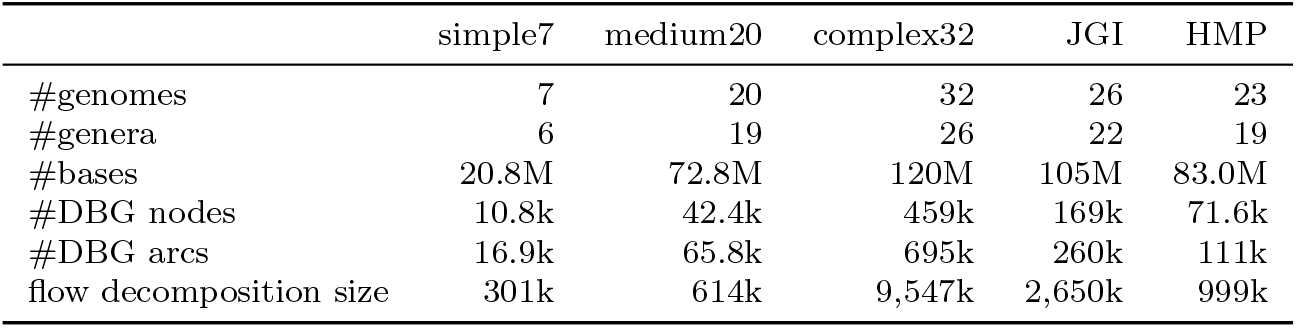
Statistics of the used datasets. The DBG is an arc-centric de Bruijn graph of order *k* = 31. The flow decomposition is that computed by our enumeration algorithm.

### Comparing assembly contiguity

In the EA50 and EA75 rows we also show in parentheses the relative improvement with respect to unitigs. In this way we can see that flowtigs give a notable improvement in comparison to structural contigs.

As we can see in Table 2, flowtigs provide consistently better EA50 and EA75 values than unitigs on all datasets (13%–17.1% better on EA50 and 13.5%–99.6% better on EA75). Moreover, our improvements over unitigs are larger on shorter contigs (EA75) than longer contigs (EA50), with the exception of HMP where these values are still very close. On the other hand, structural contigs provide only minimal improvements over unitigs: up to 2.18% for EA50 on all datasets, and up to 4.6% for EA75, on all datasets, excluding complex32. On complex32, under EA75 all contigs provide significant improvements over unitigs, which might be due to the fact that the EA75 value of unitigs is already very small (e.g. compared to the other four datasets). That is, the graph of complex32 has mostly short unitigs (i.e., to cover 75% of the genomic position one needs unitigs of length at least 538) due to branching arising from the many species in the graph; extended contigs, structural contigs and flowtigs are thus likely to be extended over such branching nodes. Furthermore, the improvement that flowtigs bring over unitigs is consistently several times higher (between 4× for EA75 in complex32 and 14× for EA50 in HMP) than what structural contigs provide over unitigs. Exceptionally, in the simple7 dataset we have an improvement of 325×, which is due to the fact that the improvement of structural contigs over unitigs is of just 0.0366%.

**Table 2:** Assembly contiguity and computation time of various tigs. The EA50 and EA75 metrics are an extension of the NGA50 and NGA75 metrics that are robust against overlapping and repeated contigs; in parentheses, we give the improvement of the respective contigs over unitigs. “gen. frac.” is the percentage of the metagenomic reference that is covered by the contigs. Since we work with perfect data, this should always be exactly 100%. But since QUAST uses approximate alignment, it is not. See Appendix A.1 for more details. “k” is the SI multiplier by 1, 000. Unitigs are computed with BCALM2 [6]. All other algorithms take unitigs as input, but for readability we do not include the computation of unitigs in their performance metrics. The computation of flowtigs contains also the run of a separate tool that transforms the node-centric DBG output by BCALM2 into an arc-centric one, since the flowtigs tool cannot directly read the output of BCALM2.

| dataset | metric | unitigs | extended contigs<br>+ unitigs | structural contigs<br>+ unitigs | flowtigs<br>+ unitigs |
| --- | --- | --- | --- | --- | --- |
| simple7 | EA50 | 21.8k | 21.8k (0.0366%) | 21.8k (0.0366%) | 24.7k (13.0%) |
|  | EA75 | 10.5k | 10.8k (2.38%) | 10.8k (2.38%) | 12.2k (16.0%) |
|  | gen. frac. | 99.75% | 99.99% | 99.99% | 99.99% |
|  | time (s) | 85.5 | 3.08 | 7.52 | 5.32 |
| medium20 | EA50 | 22.5k | 22.7k (1.05%) | 22.7k (1.05%) | 25.6k (13.5%) |
|  | EA75 | 10.4k | 10.7k (3.22%) | 10.7k (3.22%) | 12.3k (18.9%) |
|  | gen. frac. | 99.57% | 99.97% | 99.97% | 99.98% |
|  | time (s) | 200 | 11.0 | 81.0 | 17.2 |
| complex32 | EA50 | 11.1k | 11.3k (1.81%) | 11.3k (1.81%) | 12.5k (13.0%) |
|  | EA75 | 538 | 664 (23.4%) | 664 (23.4%) | 1.07k (99.6%) |
|  | gen. frac. | 94.27% | 99.86% | 99.86% | 99.79% |
|  | time (s) | 267 | 20.4 | 28,700 | 164 |
| JGI | EA50 | 16.5k | 16.8k (2.18%) | 16.8k (2.18%) | 19.3k (17.1%) |
|  | EA75 | 4.67k | 4.89k (4.6%) | 4.89k (4.6%) | 5.89k (26.0%) |
|  | gen. frac. | 99.16% | 99.96% | 99.96% | 99.97% |
|  | time (s) | 348 | 17.1 | 5,740 | 41.8 |
| HMP | EA50 | 19.4k | 19.6k (0.952%) | 19.6k (0.952%) | 22.2k (14.3%) |
|  | EA75 | 8.62k | 8.79k (2.03%) | 8.79k (2.03%) | 9.79k (13.5%) |
|  | gen. frac. | 99.23% | 99.91% | 99.91% | 99.95% |
|  | time (s) | 211 | 12.2 | 216 | 20.2 |

In Table 2 we also include the genome fraction in our table even though in theory it should be 100% in each case since we add unitigs and work on error-free data. In practice it is slightly lower, since QUAST uses approximate alignment that does not align very short contigs. If it was much lower than 100%, this may be an indicator that our EA*x* statistics are skewed, as they are based on alignments. However, it is above 99% in all but one cases and only very short contigs are unaligned, specifically all unaligned contigs are shorter than 150bp in all datasets. For the unitigs of complex32, we can assume that they contain many very short contigs, as the graph is very complex, which explains why the genome fraction is only 94% in this case. The EA*x* metrics are based on the longest contigs, similar to the well-known NGA*x* metrics. Hence, we take the genome fraction as an indicator that our statistics are accurate.

To sum up, assuming perfect data as in our experiments, flowtigs provide the first notable improvement over both long unitigs (EA50) and short unitigs (EA75). At the same time, they admit a simpler characterisation than structural contigs (i.e. excess flow). Moreover, since the definition the excess flow of a walk is independent of the flow conservation assumption, is also more amenable as a heuristic on real assembly graphs.

### Comparing performance

We also compare the performance of the various algorithms in Table 2. Note that unitigs are computed with BCALM2 [6], a highly engineered parallel external memory algorithm, while all other algorithms run single-threaded with a less engineered proof-of-concept implementation. Further, unitigs are the input to all other algorithms, but we report the runtimes of the other algorithms without that of unitigs. In a practical assembler, the graph will likely be computed with a different tool than BCALM2, hence we get a better comparison between the algorithms if we ignore the initial graph-building step.

The algorithm for extended contigs is very simple and runs in linear time, which is visible in its great performance compared to structural contigs and flowtigs. Structural contigs are computed with an *O*(*mn*) algorithm, which becomes notable on the larger graphs of complex32 and JGI, where the runtime is significantly higher than that of the other algorithms. But while flowtigs are computed with an *O*(||D_*f*_||) algorithm which is *O*(*mn*) in the worst case as well, the actual size of the flow decomposition in our graphs is much lower than that as shown in Table 1. Hence, it runs much faster and behaves more like the linear-time extended contig algorithm than the quadratic structural contig algorithm. See Appendix A.2 for an evaluation of the memory consumption of the algorithms.

## 5 Discussion and future work

In this paper we introduced flowtigs (as generalising the safe walks in DAGs from [21]) and compared them to previous safe contigs. On all the analysed datasets, flowtigs are the only safe contigs that exhibit non-negligible improvements over unitigs. For the first time in general graphs, we hope to have a notion of safe walks enjoying the same and in parts even better favourable robustness properties as unitigs and extended contigs, while potentially resulting in a much larger improvement in assembly contiguity.

In the future, we hope that we can obtain even longer safe walks than flowtigs by making use of more information that is usually available during a modern assembly process. Flowtigs use all information that is available through the structure of the assembly graph and the abundance values on the arcs. Hence, based just on this information, no further improvement can be obtained. However, in a modern assembly process, there is typically more information available that we have so far ignored. For example, in the most recent assembly of the human genome [36], an assembly graph was constructed using shorter but very accurate reads (PacBio HiFi) and then longer reads (Oxford Nanopore) were aligned to that graph. These longer reads were used to obtain very long contigs by heuristically bridging through even complex tangles in the assembly graph. Having proven their potential, it would be very interesting to see what information can safely be extracted from them in a theoretically sound model, e.g. by using them as subpath constraints for a flow decomposition. This could also result in an assembly pipeline that completely automates genome assembly such as that of Rautiainen et al. [41], but by using only safe algorithms. Such a pipeline would have the desirably property that it produces no errors given error-free data.

## A Enumerating all maximal flowtigs

### Overview

The maximal flowtigs can be identified in *O*(*mn*) time by first computing a decomposition 𝒟 of (*G, f*) into closed walks and verifying the safety of its subwalks using a two-pointer approach together with our excess flow characterisation. This is correct by definition, since safe walks are subwalks of some closed walk in any flow decomposition.

For the decomposition step, we iteratively find a closed path *P* (which is also a closed walk) and add (*P*, min_*e∈P*_ *f* (*e*)) to our decomposition until (*G, f*) is decomposed. This works in *O*(*mn*) time and space, since in each iteration we fully decompose at least one out of the *m* arcs of (*G, f*) and each closed path uses *O*(*n*) vertices.

During the two-pointer step, a recurrent operation is that of updating the excess flow of a walk when extending it in the end or the beginning. For that we present a lemma originally presented in Khan et al. [21] for DAGs, which also holds for our case as excess flow does not depend on the type of graph in question.

#### Lemma 12 ([21])

*For any walk in a flow graph* (*G, f*), *adding an arc e* = (*u, ν*) *to its start or its end, reduces its excess flow by f*_*in*_(*ν*) *− f* (*e*), *or f*_*out*_(*u*) *− f* (*e*), *respectively. Analogously, removing the arc e* = (*u, ν*) *from its start or its end, increases its excess flow by f*_*in*_(*ν*) *− f* (*e*), *or f*_*out*_(*u*) *− f* (*e*), *respectively. The quantities f*_*in*_(*ν*) *and f*_*out*_(*ν*) *can be computed in O*(*m* + *n*) *time*.

### Two-pointer phase

Based on a decomposition of (*G, f*), we can compute the maximal flowtigs along each closed walk *D* ∈ 𝒟 using a two-pointer algorithm as follows. We start with the subwalk containing the first arc of *D*. We compute its excess flow *x*, and while *x >* 0 we append the next arc to the walk on the right and incrementally compute its excess flow by Lemma 12. Whenever *x ≤* 0, we store the flowtig between the left pointer and the arc preceding the right pointer, and we move the left pointer forward, effectively removing the first arc of the walk, and update the excess flow similarly by Lemma 12. Note that these updates take constant time. We stop when the left pointer returns to the first arc of *D*, implying that we have tried every possible arc in *D* to be the beginning of a flowtig.

### Ensuring maximality

Note that these flowtigs may only be maximal with respect to the closed walk from where they were scanned. That is, it may happen that a flowtig can be extended using arcs from another closed walk of the decomposition. In any case, to ensure maximality, we check if the walk can be extended on the right and then on the left using arcs of maximum flow. Extending a walk with an arc of maximum flow maximises the excess flow of the extended walk, and thus the unsafety of such an extension implies the unsafety of all other extensions. Therefore, if any extension (left or right) preserves safety, then the walk is not maximal safe since we can make it longer. On the other hand, if both extensions individually render the walk unsafe, then the walk is maximal safe. This check can be done in constant time by precomputing a flow-maximum incoming and outgoing arc of every node, which can be done in *O*(*m* + *n*).

### Handling recurring cycles

There is an important detail that must be handled in order to achieve the claimed *O*(*mn*) runtime, which is in computing safe recurring cycles, i.e. walks along closed walks that repeat arcs as long as their excess flow is positive (note that in Figure 2b we can make the flow values of the inner square arbitrarily large while preserving flow conservation in every node, e.g. by adding a value *k* ∈ℚ^+^ to every arc of the square. In turn, this would make the length of the flowtig drawn in green arbitrarily large). Thus, applying a standard two pointer algorithm would yield an algorithm with only pseudo-polynomial runtime, as the length of such walks depend on the flow values of the arcs. To avoid this, we use the following lemma, together with an argument based on flow conservation.

#### Lemma 13.

*A safe recurring cycle W with respect to a cycle C* = (*ν*_1_, *e*_1_, …, *ν*_*ℓ*_, *e*_*ℓ*_, *ν*_1_) *crosses from e*_*ℓ*_ *to e*_1_ *at most* 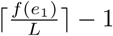 *times, where L denotes the leakage of the walk Ce*_1_.

*Proof*. Suppose that the recurring walk *W* consists of *c >* 0 repetitions of the cycle *C*, followed by a non-empty prefix of *C*. If *W* is a cycle, then it does not cross from *e*_*k*_ to *e*_1_. Otherwise, every time *W* walks over *Ce*_1_ it leaks exactly *L* flow, and so it would leak *c* · *L* when doing so *c* times. Since *W* is safe, we have 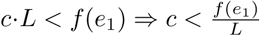. Thus, *c* is at most the greatest integer strictly smaller than, 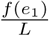, i.e. 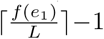. □

Now we will argue that bootstrapping the two pointer phase with Lemma 13 in the first scanned arc of *D* ∈ *𝒟* is enough to achieve the desired runtime. Clearly, the left pointer moves *n* times, so it remains to analyse the behaviour of the right pointer.

Note that while the left pointer moves from the first to the last arc of *D* it decreases the leakage by at most *L*. Now, if *f*_*M*_ and *f*_*m*_ denote the maximum and minimum flow occurring in the arcs of *D*, then *f*_*M*_ *− f*_*m*_ *≤ L*, since to get from *f*_*M*_ to *f*_*m*_ we need to leak at least *L*, otherwise *f*_*m*_ would not be the minimum flow value occurring in *D*. Then, when the left pointer reaches the last arc of *D* it has introduced at most 2 *L* of flow to be consumed by the right pointer. Since the right pointer consumes *L* units of flow per complete round around *D* (plus the next arc) and since the computation of the suffix of the first flowtig uses in the worst case *n −* 1 arcs (after the initial “jump”) we conclude that the right pointer does *O*(*n*) transitions in total.

### Storing walks efficiently

A similar problem to that of computing safe recurring cycles is in their *representation* since we can not afford to store them explicitly as sequences of arcs. Instead, we use a compact representation inspired by Khan et al. [21]. Let *𝒟* = {*D*_1_, …, *D*_*k*_} be a flow decomposition and let *W* be a flowtig. Then, we represent *W* by three integers (*i, j, l*), where *i* points to the closed walk *D*_*i*_ ∈ *𝒟* where *W* occurs, *j* points to the first arc of *W* in *D*_*i*_, and *l* denotes the length of *W* . This encoding allows us to uniquely identify flowtigs with additional constant space per flowtig.

### Optimal enumeration of flowtigs

From the discussion above, we conclude that we can identify every maximal flowtig in an instance (*G, f*) of the flow decomposition problem. We summarise this in the following theorem.

#### Theorem 2 (Optimal enumeration of flowtigs)

In Figure 4 we present a family of graphs requiring *Ω*(*mn*) total space to decompose its flow and containing Θ(*mn*) distinct maximal flowtigs. This structure is identical to the one presented in [21], except that we add an arc from *b*_*k*_ to *a*_1_ to make the graph into a circulation.

**Figure 4:**
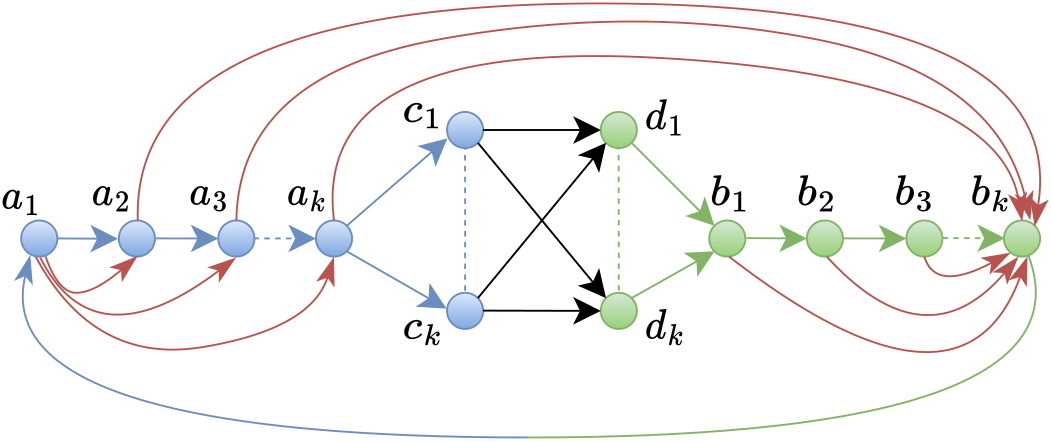
A family of graphs parameterized by *k* ≥ 1 with Θ(*k*) nodes, at least Θ(*k*) arcs (sparsest case) and at most Θ(*k*^2^) arcs (densest case). The black arcs have *k* flow, the red arcs have unit flow, and the remaining arcs are according to flow conservation. Such flow graphs contain Θ(*mn*) unique maximal flowtigs, showing that our algorithm is optimal.

About the structure of the graph, observe that by choosing *k* = *n/*4 and any subset of connections between *C* = {*c*_1_, …, *c*_*k*_} and *D* = {*d*_1_, …, *d*_*k*_} that make the graph strongly connected, we get a graph with any *n* and *m*. Let there be flow *k* on the black arcs and unit flow on the red arcs. The remaining arcs are according to flow conservation. Further, note that *m* = Θ(|*C* × *D*|) and that *n* = Θ(*k*).

For the analysis of the flow decomposition *𝒟*, note that each arc in *C* × *D* necessarily contributes in *𝒟* with at least one closed walk of length *at least* Θ(*k*), implying that ||𝒟|| = *Ω*(*mn*).

For the analysis of the maximal flowtigs, observe that there are |*C* × *D*| distinct flowtigs from *a*_*i*_ to *b*_*i*_ for all 1 *≤ i ≤ k* because every path from *a*_*i*_ to *b*_1_ has excess flow *i*. For all 2 *≤ j ≤ k −* 1, the *a*_*j*_-*b*_*j*_ flowtigs are maximal, and the *a*_1_-*b*_1_ and *a*_*k*_-*b*_*k*_ flowtigs can only be extended by adding the green and blue arc to the beginning and end, respectively. Therefore, we have *Ω*(|*C* × *D*| · *k*) = *Ω*(*mn*) distinct maximal flowtigs.

To conclude our analysis, recall that for any flow graph there is a *O*(*mn*)-sized flow decomposition, and so the graph shown in Figure 4 only admits flow decompositions of size Θ(*mn*). Moreover, the *number* of distinct maximal flowtigs is bounded above by the size of the smallest flow decomposition in the underlying graph. Indeed, this is true by definition of safety, and due to the fact that in any closed walk *D* ∈ *𝒟* we have at most one unique maximal flowtig per arc in *D*. Then, Figure 4 has Θ(*mn*) distinct maximal flowtigs, as we wanted. This shows that our algorithm is optimal in time and space.

### Deduplication

Finally, our algorithm may produce duplicate maximal flowtigs. To address this we use a suffix tree with suffix links to identify the set of maximal flowtigs without duplicates, similar to Obscura et al. [37]. This increases the runtime of our algorithm to *O*(||*𝒟*_*f*_ || log *m*).

A maximal flowtig may be reported multiple times when there exist distinct *D*_*i*_, *D*_*j*_ ∈ *𝒟* such that *D*_*i*_ ∩ *D*_*j*_ ≠ *∅*. However, maximal flowtigs that repeat any arc cannot be reported multiple times, as that would imply that there exist two equivalent cycles in the decomposition, which can not happen in our algorithm. Hence, we only need to deduplicate paths and cycles.

For filtering out duplicates efficiently, we can apply an idea similar to that of Obscura et al. [37]. We build a string *S* := *D*_1_*D*_1_*D*_1_*D*_1_*D*_1_*D*_1_#*D*_2_*D*_2_*D*_2_*D*_2_*D*_2_*D*_2_# … #*D*_*k*_*D*_*k*_*D*_*k*_*D*_*k*_*D*_*k*_*D*_*k*_ of length *O*(||*𝒟*_*f*_ ||) for our decomposition {*D*_1_, …, *D*_*k*_} which contains the arcs of our decomposing cycles. Then we build a suffix tree in *O*(||*𝒟*_*f*_||) time using an algorithm by Farach [7], which we can apply since our nodes form an integer alphabet. Then we add suffix links into the tree using a linear-time algorithm by Maaß [28]. Suffix links are arcs that point from a string *a*_1_ … *a*_*ℓ*_ to a string *a*_2_ … *a*_*ℓ*_.

Now, we can walk along the suffix tree while executing the two-pointer algorithm. After finding the initial walk in the two-pointer algorithm, we check if the initial walk completes the decomposing cycle at least twice. If it does, then all maximal flowtigs reported by the two-pointer algorithm will repeat at least one arc, and hence cannot have duplicates. In this case, we do not need to use the suffix tree.

If on the other hand the initial walk completes the decomposing cycle less than two times, we make use of the suffix tree. We first traverse the suffix tree from the root to find the initial walk, and then traverse one step deeper for each move of the right pointer, and use a suffix link for each move of the left pointer. Since the right pointer moves at most three times around the cycle, and the initial walk repeats the cycle less than two times, repeating each decomposing cycle six times in the string *S* is enough for never entering a leaf when moving the right pointer.

Using a suffix link takes constant time. However, when stepping deeper into the tree, we have up to *O*(*m*) options, of which only is correct. To choose the right one we can use binary search which takes *O*(log *m*) time. Hence, while executing the two-pointer algorithm, we can traverse the suffix tree in constant time for each left pointer update, and in *O*(log *m*) for each right pointer update. In total this takes *O*(||*𝒟*_*f*_|| log *m*) time for all executions of the two-pointer algorithm.

Using the simultaneous traversal of the suffix tree, we can mark nodes of the suffix tree whenever we output a maximal flowtig. Then, before we output a maximal flowtig, we check if the corresponding node was marked already, and only output it if the node is unmarked.

### A.1 Implementation and evaluation

We implement the flowtig algorithm in Rust. Instead of the complex deduplication algorithm we use a hash set to deduplicate the output. The implementation is available on github [13] and on Software Heritage [14] under the BSD-2-Clause license. Our experiment pipeline is written with snakemake [33] and available on Software Heritage [16] under the BSD-2-Clause license. To compare against extended and structural contigs, we use the implementation by Schmidt available on Software Heritage [43] under the BSD-2-Clause license.

The algorithms for extended contigs and structural contigs use a compact representation of the input strings, storing all strings 2-bit encoded in a single bitvector. Further, they never make any copies of these strings, and hence use only a low amount of memory. Compared to that, our flowtig implementation stores strings in ASCII format. During deduplication, it copies the unitigs into the flowtigs and stores them inside a hash set, hence it holds the input strings and all flowtigs simultaneously in memory. This causes it to use around an order of magnitude more memory. If we were to use the compact string format for flowtigs as well and would deduplicate while storing the flowtigs as sequences of arcs without copying the strings, then our memory usage would likely be similar to that of structural contigs.

When computing longer safe walks, we get an issue with a lower genome fraction that was already noticed by Jain [19] on overlap graphs. By only using maximal contigs, some genomes may be left with a gap where no contig aligns, even on perfect data. See for example Figure 1, where in (c) the middle horizontal arc is not covered by any contig that aligns to the red genome, and in (d) there is no contig that aligns to the blue genome outside of the repeat. To mitigate this effect, we always run our evaluations on the maximal contigs combined with unitigs. For example with flowtigs on complex32, the genome fraction would only be 99.29% without adding unitigs, while it is 99.77% with adding unitigs. See Table 3 for the genome fractions of flowtigs on all datasets with and without adding unitigs.

**Table 3:** Genome fraction of flowtigs without and with adding unitigs.

| Genome | only flowtigs | flowtigs + unitigs |
| --- | --- | --- |
| simple7 | 99.96 | 99.99 |
| medium20 | 99.96 | 99.98 |
| complex32 | 99.29 | 99.79 |
| JGI | 99.94 | 99.97 |
| HMP | 99.92 | 99.95 |

We compare flowtigs against unitigs, extended contigs, and structural contigs. For each metagenomic reference, we circularise each genome, and then compute a compacted de Bruijn graph for with *k*-mer abundances for *k* = 31 using bcalm2 [6]. Since bcalm2 does not support setting a multiplier for each genome, we simply feed bcalm2 *a* copies for a genome with abundance *a*. The sizes of the de Bruijn graphs are displayed in Table 1. From the compacted de Bruijn graph we then compute the various tigs. We evaluate the tigs with QUAST [31] using various common metrics. Since we use error-free data, we do not use metaQUAST [32], because its changes over QUAST are only regarding erroneous data. In addition to QUAST’s metrics, because flowtigs can overlap a lot, we add the EA50 family of metrics, which is an improvement over NGA50 that is robust against overlapping contigs [44]. The EA50 is computed by aligning the contigs to the reference, and for each reference base identifying the longest contig that aligns to it. These lengths are then sorted and for e.g. EA50, the 50-percentile is reported. EA75 works analogously, by reporting the 75-percentile of largest values. We implement this metric in a modified version of QUAST which additionally computes average contig lengths and does not filter out short contigs. It is available on Software Heritage [17] under QUAST’s original custom license.

Note that QUAST filters alignments below a length of 65 regardless of its parameterisation, hence very short contigs do not align in our evaluation, and we get a genome fraction below 100% even though we work with perfect data and safe algorithms only.

### A.2 Comparing memory usage

We compare the performance of the various algorithms including also memory usage in Table 4. We see that extended contigs and structural contigs are very similar, while flowtigs use roughly an order of magnitude more memory, up to almost 4GiB. This is because extended contigs and structural contigs use compact data structures to store strings and do not require deduplication, while our flowtigs implementation uses simpler methods to store strings as well as a simple method for deduplication. We assume that using a more compact method to store strings and an optimised way to do deduplication would reduce the RAM usage of flowtigs down to around that of structural contigs. See Appendix A.1 for details.

**Table 4:**
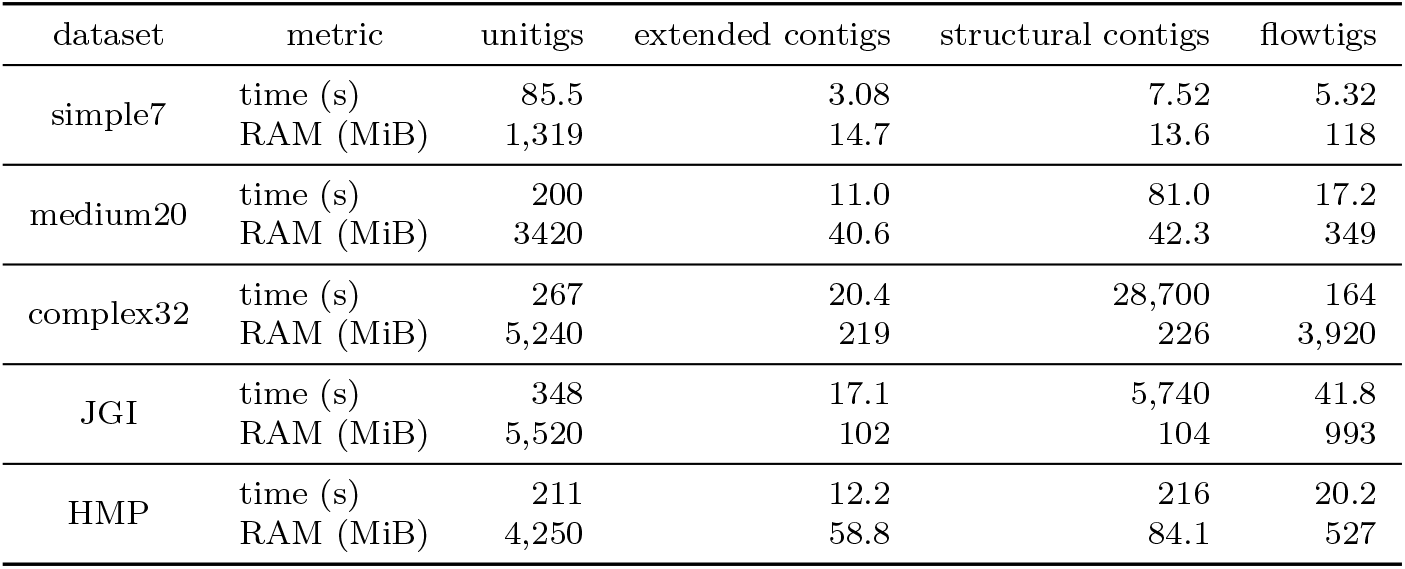
Performance for computing various tigs. Values are rounded to the three most significant digits. Unitigs are computed by BCALM2 [6] which uses external memory and runs in parallel. All other algorithms take unitigs as input, but for readability we do not include the computation of unitigs in their performance metrics. The computation of flowtigs contains also the run of a separate tool that transforms the node-centric DBG output by BCALM2 into an arc-centric one, since the flowtigs tool cannot directly read the output of BCALM2.

| dataset | metric | unitigs | extended contigs | structural contigs | flowtigs |
| --- | --- | --- | --- | --- | --- |
| simple7 | time (s) | 85.5 | 3.08 | 7.52 | 5.32 |
|  | RAM (MiB) | 1,319 | 14.7 | 13.6 | 118 |
| medium20 | time (s) | 200 | 11.0 | 81.0 | 17.2 |
|  | RAM (MiB) | 3420 | 40.6 | 42.3 | 349 |
| complex32 | time (s) | 267 | 20.4 | 28,700 | 164 |
|  | RAM (MiB) | 5,240 | 219 | 226 | 3,920 |
| JGI | time (s) | 348 | 17.1 | 5,740 | 41.8 |
|  | RAM (MiB) | 5,520 | 102 | 104 | 993 |
| HMP | time (s) | 211 | 12.2 | 216 | 20.2 |
|  | RAM (MiB) | 4,250 | 58.8 | 84.1 | 527 |

## References

[1] Ravindra K. Ahuja, Thomas L. Magnanti, and James B. Orlin. Network Flows: Theory, Algorithms, and Applications. USA: Prentice-Hall, Inc., 1993. isbn: 013617549X.

[2] Martin Ayling, Matthew D Clark, and Richard M Leggett. “New approaches for metagenome assembly with short reads”. In: Briefings in Bioinformatics 21.2 (Feb. 2019), pp. 584–594. issn: 1477-4054. doi: 10.1093/bib/bbz020. eprint: https://academic.oup.com/bib/article-pdf/21/2/584/33583908/bbz020.pdf.

[3] Gaëtan Benoit, Sébastien Raguideau, Robert James, Adam M. Phillippy, Rayan Chikhi, and Christopher Quince. “Efficient High-Quality Metagenome Assembly from Long Accurate Reads using Minimizer-space de Bruijn Graphs”. In: bioRxiv (2023). doi: 10.1101/2023.07.07.548136. eprint: https://www.biorxiv.org/content/early/2023/07/08/2023.07.07.548136.full.pdf.

[4] Manuel Cáceres, Brendan Mumey, Edin Husić, Romeo Rizzi, Massimo Cairo, Kristoffer Sahlin, and Alexandru I. Tomescu. “Safety in multi-assembly via paths appearing in all path covers of a DAG”. In: IEEE/ACM Transactions on Computational Biology and Bioinformatics 19.6 (2022), pp. 3673–3684. url: 10.1109/TCBB.2021.3131203.

[5] Massimo Cairo, Shahbaz Khan, Romeo Rizzi, Sebastian Schmidt, Alexandru I Tomescu, and Elia C Zirondelli. “Cut Paths and Their Remainder Structure, with Applications”. In: 40th International Symposium on Theoretical Aspects of Computer Science (STACS 2023). Schloss Dagstuhl-Leibniz-Zentrum für Informatik. 2023.

[6] Rayan Chikhi, Antoine Limasset, and Paul Medvedev. “Compacting de Bruijn graphs from sequencing data quickly and in low memory”. In: Bioinformatics 32.12 (2016), pp. i201–i208.

[7] Martin Farach. “Optimal suffix tree construction with large alphabets”. In: Proceedings 38th Annual Symposium on Foundations of Computer Science. IEEE. 1997, pp. 137–143.

[8] Xiaowen Feng, Haoyu Cheng, Daniel Portik, and Heng Li. “Metagenome assembly of high-fidelity long reads with hifiasm-meta”. In: Nature Methods 19.6 (2022), pp. 671–674.

[9] Adrian Fritz, Peter Hofmann, Stephan Majda, Eik Dahms, Johannes Dröge, Jessika Fiedler, Till R Lesker, Peter Belmann, Matthew Z DeMaere, Aaron E Darling, et al. “CAMISIM: simulating metagenomes and microbial communities”. In: Microbiome 7.1 (2019), pp. 1–12.

[10] Hadrien Gourlé, Oskar Karlsson-Lindsjö, Juliette Hayer, and Erik Bongcam-Rudloff. “Simulating Illumina metagenomic data with InSilicoSeq”. In: Bioinformatics 35.3 (2019), pp. 521–522.

[11] Steffen Heber, Max Alekseyev, Sing-Hoi Sze, Haixu Tang, and Pavel A Pevzner. “Splicing graphs and EST assembly problem”. In: Bioinformatics 18.suppl 1 (2002), S181–S188.

[12] Ramana M Idury and Michael S Waterman. “A new algorithm for DNA sequence assembly”. In: Journal of Computational Biology 2.2 (1995), pp. 291–306.

[13] Eliel Ingervo. Flowtigs. Version 1.0.1. Oct. 2023. url: https://github.com/elieling/flowtigs.

[14] [SW] Eliel Ingervo, Flowtigs 2023. swhid: ⟨swh:1:rev:2685085eab02c124b8a62787bf75e4922b252882;origin=https://github.com/elieling/flowtigs;visit=swh:1:snp:924fdd2f176a8c0c2f0498debb423ab4e33ea7f7⟩.

[15] Eliel Ingervo. Flowtigs datasets. Version v1. 2023. doi: 10.5281/zenodo.8434267.

[16] [SW] Eliel Ingervo, Flowtigs experiment pipeline 2023. swhid: ⟨swh:1:rev:c5db004c628c665c0cd4043a0550011d0502c67f;origin=https://github.com/elieling/safe-paths-with-flowtigs;visit=swh:1:snp:4f699a2bd0ea9ec8740492c6e77956bbcb426ecb⟩.

[17] [SW] Eliel Ingervo and Sebastian Schmidt, Quast 2023. swhid: ⟨swh:1:rev:cf3870b84449d69de76cbb704f989c433a34e6f0;origin=https://github.com/elieling/quast;visit=swh:1:snp:7e09bcde042226387c692410a0fbbc3dd8c06332⟩.

[18] Benjamin Grant Jackson. Parallel methods for short read assembly. Iowa State University, Ph.D. thesis, 2009.

[19] Chirag Jain. “Coverage-preserving sparsification of overlap graphs for long-read assembly”. In: Bioinformatics 39.3 (2023), btad124.

[20] Evgeny Kapun and Fedor Tsarev. “De Bruijn Superwalk with Multiplicities Problem is NP-hard”. In: BMC Bioinformatics 14 Supplement 5.S7 (2013), pp. 1–4. doi: 10.1186/1471-2105-14-S5-S7.

[21] Shahbaz Khan, Milla Kortelainen, Manuel Cáceres, Lucia Williams, and Alexandru I. Tomescu. “Safety and Completeness in Flow Decompositions for RNA Assembly”. In: RECOMB 2022 - 26th Annual International Conference on Research in Computational Molecular Biology. Vol. 13278. Lecture Notes in Computer Science. Springer, 2022, pp. 177–192. doi: 10.1007/978-3-031-04749-7_11.

[22] Carl Kingsford, Michael C. Schatz, and Mihai Pop. “Assembly complexity of prokaryotic genomes using short reads”. In: BMC Bioinformatics 11.21 (2010), pp. 1–11. doi: 10.1186/1471-2105-11-21.

[23] Mikhail Kolmogorov, Derek M Bickhart, Bahar Behsaz, Alexey Gurevich, Mikhail Rayko, Sung Bong Shin, Kristen Kuhn, Jeffrey Yuan, Evgeny Polevikov, Timothy PL Smith, et al. “metaFlye: scalable long-read metagenome assembly using repeat graphs”. In: Nature Methods 17.11 (2020), pp. 1103–1110.

[24] Mikhail Kolmogorov, Jeffrey Yuan, Yu Lin, and Pavel A Pevzner. “Assembly of long, error-prone reads using repeat graphs”. In: Nature Biotechnology 37.5 (2019), pp. 540–546.

[25] Dinghua Li, Chi-Man Liu, Ruibang Luo, Kunihiko Sadakane, and Tak-Wah Lam. “MEGAHIT: an ultra-fast single-node solution for large and complex metagenomics assembly via succinct de Bruijn graph”. In: Bioinformatics 31.10 (2015), pp. 1674–1676.

[26] Wei Li. RNASeqReadSimulator: a simple RNA-seq read simulator. 2014. url: http://alumni.cs.ucr.edu/~liw/rnaseqreadsimulator.html.

[27] Lei Liu, Yulin Wang, You Che, Yiqiang Chen, Yu Xia, Ruibang Luo, Suk Hang Cheng, Chunmiao Zheng, and Tong Zhang. “High-quality bacterial genomes of a partial-nitritation/anammox system by an iterative hybrid assembly method”. In: Microbiome 8 (2020), pp. 1–17.

[28] Moritz G Maaß. “Computing suffix links for suffix trees and arrays”. In: Information Processing Letters 101.6 (2007), pp. 250–254.

[29] Veli Mäkinen, Djamal Belazzougui, Fabio Cunial, and Alexandru I Tomescu. Genome-Scale Algorithm Design: Bioinformatics in the Era of High-Throughput Sequencing. Cambridge University Press, 2023.

[30] Paul Medvedev, Konstantinos Georgiou, Gene Myers, and Michael Brudno. “Computability of models for sequence assembly”. In: International workshop on algorithms in bioinformatics. Springer. 2007, pp. 289–301.

[31] Alla Mikheenko, Andrey Prjibelski, Vladislav Saveliev, Dmitry Antipov, and Alexey Gurevich. “Versatile genome assembly evaluation with QUAST-LG”. In: Bioinformatics 34.13 (2018), pp. i142–i150.

[32] Alla Mikheenko, Vladislav Saveliev, and Alexey Gurevich. “MetaQUAST: evaluation of metagenome assemblies”. In: Bioinformatics 32.7 (2016), pp. 1088–1090.

[33] Felix Mölder, Kim Philipp Jablonski, Brice Letcher, Michael B Hall, Christopher H Tomkins-Tinch, Vanessa Sochat, Jan Forster, Soohyun Lee, Sven O Twardziok, Alexander Kanitz, et al. “Sustainable data analysis with Snakemake”. In: F1000Research 10 (2021).

[34] Eugene W Myers. “The fragment assembly string graph”. In: Bioinformatics 21.suppl 2 (2005), ii79–ii85.

[35] Sergey Nurk, Dmitry Meleshko, Anton Korobeynikov, and Pavel A Pevzner. “metaSPAdes: a new versatile metagenomic assembler”. In: Genome research 27.5 (2017), pp. 824–834.

[36] Sergey Nurk et al. “The complete sequence of a human genome”. In: Science 376.6588 (2022), pp. 44–53. doi: 10.1126/science.abj6987.

[37] Nidia Obscura Acosta, Veli Mäkinen, and Alexandru I Tomescu. “A safe and complete algorithm for metagenomic assembly”. In: Algorithms for Molecular Biology 13.3 (2018), pp. 1–12.

[38] Nidia Obscura Acosta and Alexandru I. Tomescu. “Simplicity in Eulerian circuits: Uniqueness and safety”. In: Information Processing Letters 183 (2024), p. 106421. issn: 0020-0190. doi: 10.1016/j.ipl.2023.106421.

[39] Yu Peng, Henry CM Leung, Siu-Ming Yiu, and Francis YL Chin. “Meta-IDBA: a de Novo assembler for metagenomic data”. In: Bioinformatics 27.13 (2011), pp. i94–i101.

[40] Christopher Quince, Alan W Walker, Jared T Simpson, Nicholas J Loman, and Nicola Segata. “Shotgun metagenomics, from sampling to analysis”. In: Nature Biotechnology 35.9 (Sept. 2017), pp. 833–844. issn: 1546-1696. doi: 10.1038/nbt.3935.

[41] Mikko Rautiainen, Sergey Nurk, Brian P. Walenz, Glennis A. Logsdon, David Porubsky, Arang Rhie, Evan E. Eichler, Adam M. Phillippy, and Sergey Koren. “Telomere-to-telomere assembly of diploid chromosomes with Verkko”. In: Nature Biotechnology (Feb. 2023). issn: 1546-1696. doi: 10.1038/s41587-023-01662-6. url: https://doi.org/10.1038/s41587-023-01662-6.

[42] Claudia Sala, Silvia Vitali, Enrico Giampieri, Italo Faria do Valle, Daniel Remondini, Paolo Garagnani, Matteo Bersanelli, Ettore Mosca, Luciano Milanesi, and Gastone Castellani. “Stochastic neutral modelling of the Gut Microbiota’s relative species abundance from next generation sequencing data”. In: BMC Bioinformatics 17.2 (Jan. 2016), S16. issn: 1471-2105. doi: 10.1186/s12859-015-0858-8.

[43] [SW] Sebastian Schmidt, practical omnitigs 2023. swhid: ⟨swh:1:rev:9fa8497c8de99a70c05474c8fa8318dce49ddeb9;origin=https://github.com/algbio/practical-omnitigs;visit=swh:1:snp:04e6ac0423d201dfab2c8d8ebe834756a6f88de9⟩.

[44] Sebastian Schmidt, Santeri Toivonen, Paul Medvedev, and Alexandru I Tomescu. “The omnitig framework can improve genome assembly contiguity in practice”. In: bioRxiv (2023). doi: 10.1101/2023.01.30.526175.

[45] Alexander Schrijver. Combinatorial optimization: polyhedra and efficiency. Vol. 24. 2. Springer, 2003.

[46] Mantas Sereika, Rasmus Hansen Kirkegaard, Søren Michael Karst, Thomas Yssing Michaelsen, Emil Aarre Sørensen, Rasmus Dam Wollenberg, and Mads Albertsen. “Oxford Nanopore R10.4 long-read sequencing enables the generation of near-finished bacterial genomes from pure cultures and metagenomes without short-read or reference polishing”. In: Nature Methods 19.7 (July 2022), pp. 823–826. issn: 1548-7105. doi: 10.1038/s41592-022-01539-7.

[47] Daria Shafranskaya, Varsha Kale, Rob Finn, Alla L. Lapidus, Anton Korobeynikov, and Andrey D. Prjibelski. “MetaGT: A pipeline for de novo assembly of metatranscriptomes with the aid of metagenomic data”. In: Frontiers in Microbiology 13 (2022). issn: 1664-302X. doi: 10.3389/fmicb.2022.981458.

[48] Esther Singer, Bill Andreopoulos, Robert M Bowers, Janey Lee, Shweta Deshpande, Jennifer Chiniquy, Doina Ciobanu, Hans-Peter Klenk, Matthew Zane, Christopher Daum, et al. “Next generation sequencing data of a defined microbial mock community”. In: Scientific data 3.1 (2016), pp. 1–8.

[49] Alexandru I Tomescu and Paul Medvedev. “Safe and complete contig assembly through omnitigs”. In: Journal of Computational Biology 24.6 (2017), pp. 590–602.

[50] Riccardo Vicedomini, Christopher Quince, Aaron E Darling, and Rayan Chikhi. “Strainberry: automated strain separation in low-complexity metagenomes using long reads”. In: Nature Communications 12.1 (2021), p. 4485.

[51] Hongyu Zheng, Cong Ma, and Carl Kingsford. “Deriving Ranges of Optimal Estimated Transcript Expression due to Nonidentifiability”. In: J. Comput. Biol. 29.2 (2022). Proceedings paper from RECOMB 2021, pp. 121–139. doi: 10.1089/cmb.2021.0444. url: https://doi.org/10.1089/cmb.2021.0444.

